# *In vivo* Feedback Control of an Antithetic Molecular-Titration Motif in *Escherichia coli* using Microfluidics

**DOI:** 10.1101/2020.02.28.952143

**Authors:** B. Shannon, C.G. Zamora-Chimal, L. Postiglione, D. Salzano, C. Grierson, L. Marucci, N.J. Savery, M. di Bernardo

**Affiliations:** DNA-Protein Interactions Unit, School of Biochemistry, University of Bristol, Bristol BS8 1TD, UK; Department of Engineering Mathematics, University of Bristol, BS8 1UB, Bristol, U.K.; Department of Electrical Engineering and Information Technology, University of Naples Federico II, 80125, Naples, Italy; School of Biological Sciences, University of Bristol, BS8 1UH, Bristol, U.K.

**Author notes:** These authors contributed equally to this work (B. Shannon, C.G. Zamora-Chimal). These authors contributed equally to this work (L. Marucci, N.J. Savery, M. di Bernardo).

## Abstract

We study both *in silico* and *in vivo* the real-time feedback control of a molecular titration motif that has been earmarked as a fundamental component of antithetic and multicellular feedback control schemes in *E. coli*. We show that an external feedback control strategy can successfully regulate the average fluorescence output of a bacterial cell population to a desired constant level in real-time. We also provide *in silico* evidence that the same strategy can be used to track a time-varying reference signal where the set-point is switched to a different value halfway through the experiment. We use the experimental data to refine and parameterize an *in silico* model of the motif that can be used as an error computation module in future embedded or multicellular control experiments.

## Introduction

Molecular titration modules have been highlighted as a fundamental component for the implementation of feedback controllers in living cells. As evaluated *in silico* in (Samaniego et al., 2016) and confirmed experimentally in (Annunziata et al., 2017), sequestration mechanisms can be used as effective error computation modules for the implementation of *in vivo* feedback control of living cells. More specifically, these modules can be used to compare some desired reference input of interest with a signal related to the phenotype one wishes to control, so as to provide an error signal that the molecular control device can use to regulate its behaviour. Molecular titration motifs are crucial for the implementation of embedded antithetic feedback controllers both *in vivo* (Aoki et al., 2019) and *in vitro* (Agrawal et al., 2019), and also for the realization of multicellular control strategies as first suggested in (Fiore et al., 2017).

The crucial role played by molecular titration motifs for feedback control of cells necessitates a more thorough study of their properties and dynamic response. In this paper we present the *in vivo* external feedback control of an engineered *E. coli* population endowed with the gene regulatory network shown in Figure 1A. The controller is implemented in microfluidics by closing the loop through an inducible promoter controlled by an antithetic *σ*/anti-*σ* module that is driven by an external reference input, see Figure 1B. An inverted fluorescence microscope senses the cell population outputs and drives a set of external actuators to change the input to the cells and so control their average fluorescence output.

**Figure 1.**
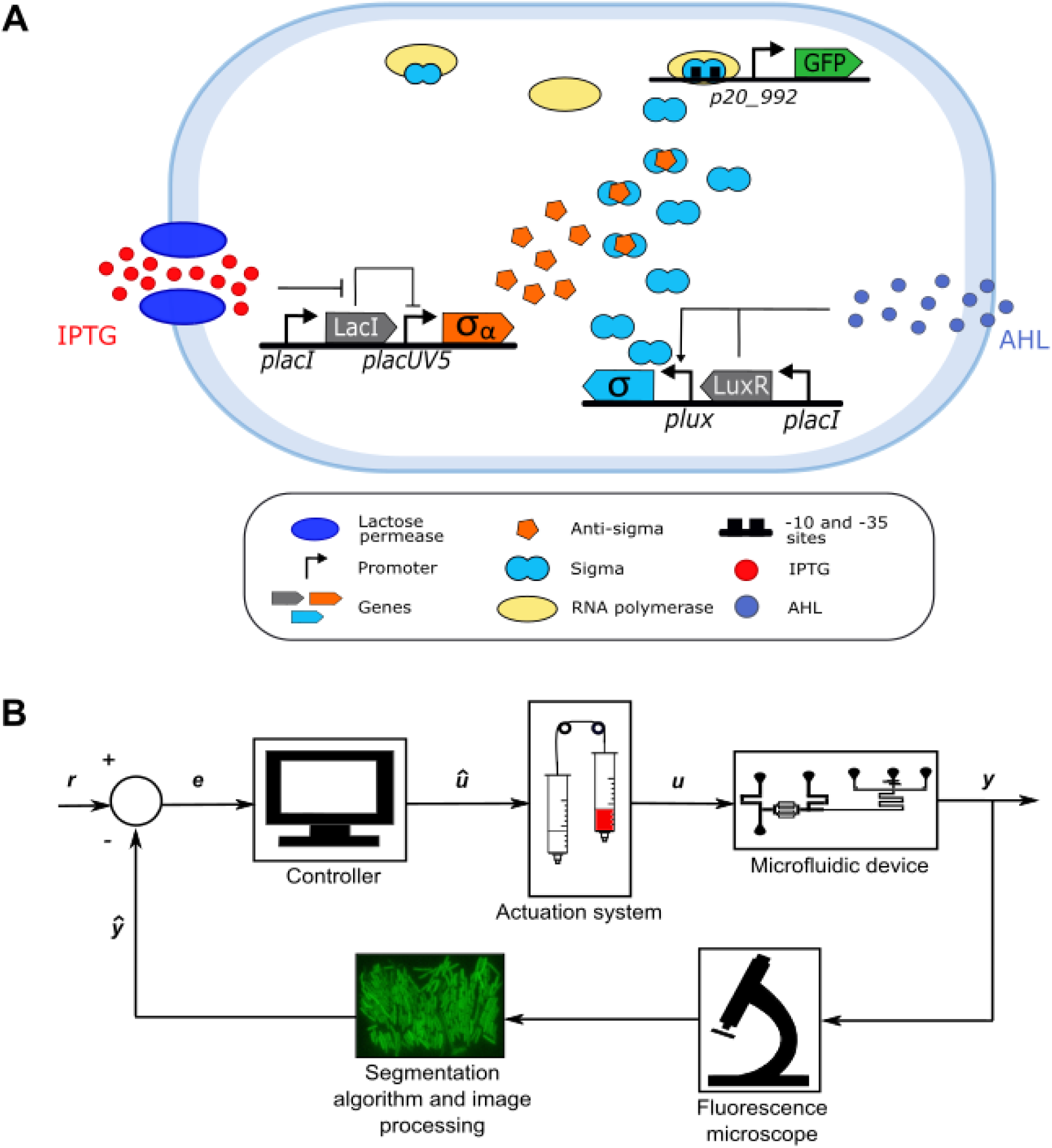
Molecular titration system within *E. coli* and external control platform. **A)** Biological design of the molecular titration system. GFP output from the system is dependent on the concentrations of IPTG and AHL. For GFP to be expressed, a specific *σ* factor is required, capable of recruiting RNA polymerase to the *p20_992* promoter upstream of GFP. The quorum sensing molecule, AHL, together with the LuxR protein, induces expression from the *plux* promoter, to produce the *σ* factor. The production of a cognate anti-*σ* factor, capable of inactivating the *σ*, occurs when the chemical inducer, IPTG, releases the Lac repressor (LacI) from the *placUV5* promoter. **B)** Schematic of the control platform. An inverted fluorescence microscope is used to capture the fluorescence (*y)* of the *E. coli* strain growing inside the microfluidic device. Images are sent to a computer running a MATLAB algorithm. This algorithm segments the images and quantifies average fluorescence (*ŷ*). The error (*e*) between the desired fluorescence value and the actual output is then evaluated and used to decide if the input should be provided to the cells or removed (*û*). The actuation system is then instructed to either deliver or remove inducers(s) from the microfluidic device accordingly.

The experiments reported here integrate and expand the preliminary investigation of the dynamic response of the module we previously explored solely through a set of batch-culture experiments in (Annunziata et al., 2017). The results we describe here are crucial to understand the ability of the module to work in real-time conditions, assessing its settling time, robustness and the quality of its response when tested in a microfluidics set-up. We also present a new *in vivo* implementation of a real-time external feedback control scheme for *E. coli* which is entirely implemented via microfluidics, the only other currently characterised example being the control of a toggle-switch presented in (Lugagne et al., 2017), which relies on the use of an entirely different microfluidics chip design, segmentation algorithm and control implementation.

## Results

### Benchmark molecular titration module

We implemented the biological system proposed and designed in (Annunziata et al., 2017)as a benchmark to test the dynamic response of an error computation module — a comparator — based on molecular titration (Rhodius et al., 2013). The system is a multi-input, single output device that uses two independent signals to regulate the expression level of a target protein of interest. It is based around an orthogonal *σ*/anti-*σ* pair able to specifically regulate the activity of the *p20_992* promoter through antithetic behaviour (Rhodius et al., 2013) (Figure 1A). The cell-cell communication molecule 3-O-C6-HSL (AHL) (Mattmann and Blackwell, 2010) and the chemical inducer Isopropyl β-D-1-thiogalactopyranoside (IPTG) were used as external inputs to control the system. The *p20_992* promoter was placed upstream of the super folder GFP (sfGFP) protein. sfGFP was chosen for its fast maturation time of approximately 6 minutes (Pedelacq et al., 2006) and for easy integration of the system into our control platform, which utilised an inverted widefield fluorescent microscope. Production of *σ*_20_992 was placed under control of the AHL inducible promoter *plux*, whilst the production of the anti-*σ* _20_992 was placed under the control of the IPTG inducible promoter *plac-UV5*. This system works on the basis that free anti-*σ* can bind to *σ*, preventing it from recruiting RNA polymerase to the *p20_992* promoter for the expression of GFP. Therefore, the level of GFP expression will be proportional to the amount of free *σ* available, and to the difference between the two signals (AHL and IPTG). All proteins were fused to a degradation tag to ensure the fast dynamics of expression from the system (Gottesman et al., 1998).

### Development of the platform for external control experiments

In order to implement external control experiments *in vivo*, we developed a microfluidics/microscopy platform for the real-time monitoring and control of our chosen *E. coli* population (see Figure 1B). The platform consisted of a microfluidic device (PDMS chip), as described in (Mondragon-Palomino et al., 2011), fixed onto the stage of an inverted widefield fluorescence microscope and connected via fluidic lines to motorized syringes for the delivery of nutrients and inducers into the chip via hydrostatic pressure. The device contained a series of trapping chambers, each allowing approximately 200 cells to grow in a monolayer. To control the level of output (fluorescence) from a monolayer population within one chamber, time-lapse imaging was performed together with real-time quantification of the average fluorescence output across cells in the chamber using an image-processing algorithm (programmed in MATLAB). Our segmentation algorithm was adapted for bacteria from (Menolascina et al., 2014) and used to segment the phase contrast images, locate the cells and obtain a binary mask, which was then used to calculate the individual cell fluorescence and compute its average across the population at every time-point. Further information on the segmentation algorithm is provided in the Methods Section.

### Open-loop system response and in silico modelling

We started by characterizing experimentally the response of the molecular titration system to time-varying external inputs. We showed that the microfluidic platform can maintain bacterial population growth, and that the GFP fluorescence of the cells can be varied as a function of the external IPTG and AHL signals. The observed experimental response is shown in Figure S1.

The experimental data was used to parameterize a computational model of the titration module (see Methods for further information). The model was taken from the equations proposed earlier in (Annunziata et al., 2017), as the following set of Ordinary Differential Equations (ODEs):

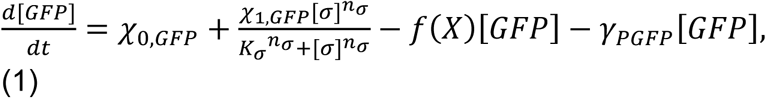

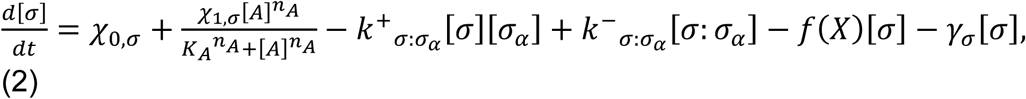

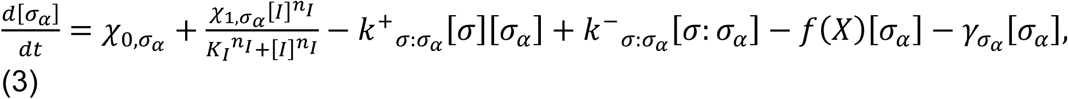

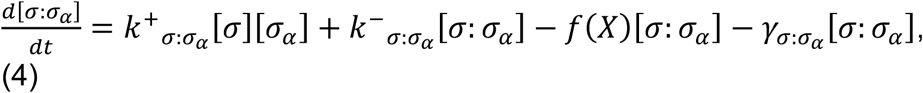

whose dependent variables model the concentrations in time of mature GFP [*GFP*], free *σ* factor [*σ*], free anti-*σ* [*σ*_*α*_] and the *σ*:anti-*σ* complex [*σ*: *σ*_*α*_]. Their dynamics are determined by production, dilution and degradation terms. In the model, the dynamics of mRNA transcription was assumed to be faster than the translation process, and a quasi-steady-state approximation of mRNA dynamics was therefore applied. Also, the dynamics of intracellular mRNAs are described through saturating Hill curves parametrised by the concentration of an input, AHL ([*A*]) or IPTG ([*I*]), or an RNA polymerase sigma subunit [*σ*], a half-maximal activation concentration (*K*_*σ*_, *K*_*A*_, and *K*_*I*_) and a Hill coefficient (*n*_*σ*_, *n*_*A*_ and *n*_*I*_). Parameters *χ*_0,*i*_ and *χ*_1,*i*_ are the basal and maximal rate of production of protein species 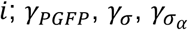 and 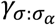 are the rates of dilution of GFP, *σ*, anti-*σ* and the σ: σ_*a*_ complex, respectively; 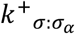 and 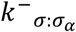 are the association and dissociation rate of the σ:σ_α_ complex formation.

In Equation (4), the function 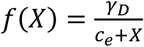 captures the competition between available ssrA tag for available proteases, where *X* = [*σ*] + [*σ*_*α*_] + [*σ*: *σ*_*α*_] + [*GFP*] is the total number of ssrA tagged proteins present in the system, *γ*_*D*_ defines the maximum value of enzymatic degradation, and *c*_*e*_ defines the half activation threshold.

Model fitting and predictions were found to match the *in vivo* observations as reported in Figure S1, confirming the viability of the computational model we derived (see Methods) for control system design and *in silico* validation.

### Closed loop control experiments

Using the *in silico* model, we designed external closed loop control experiments in which the aim was to keep GFP fluorescence at a fixed set-point or track a time-varying signal. These experiments were first performed *in silico* and then experimentally *in vivo*. AHL (10^−2^ mM) was continually supplied to the cells to maintain production of the *σ* factor, while IPTG was used as a control input (for the generation of anti-*σ* factor) by switching between 0 mM and 10^−1^ mM.

Prior to the control start in both *in silico* experiments and *in vivo* experiments, a calibration phase was performed to calculate the minimum and maximum relative fluorescence values exhibited by the cells at steady state for each of the conditions (Annunziata et al., 2017). Specifically, to calculate minimum GFP, cells were exposed to 10^−1^ mM IPTG for 300 minutes and the average GFP obtained in the last 30 minutes of this phase was normalised to 0. To calculate maximum GFP, IPTG was then removed for 300 minutes and GFP values obtained in the last 30 minutes of this phase were averaged and normalised to 1. The dynamic range of GFP was therefore set between 0 and 1 and the desired GFP reference value (*r)* to be attained was then chosen within this interval.

To implement feedback control, we used a simple yet effective relay (ON/OFF) control strategy to modulate the amount of IPTG provided to the cells as a function of the mismatch, say *e(t)*, between the refence signal (*r*) and the system fluorescence output (GFP) (Astrom and Murray, 2008). Specifically, we set

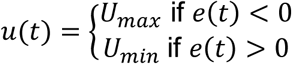

where *u(t)* is the IPTG input to cells set to *U*_*max*_ = 10^−1^ mM or *U*_*min*_ = 0mM.

To determine the feasibility of our control strategy, we first performed *in silico* stochastic simulations of the planned *in vivo* closed loop experiments. We embedded the ODE model into the agent-based simulator BSim (Matyjaszkiewicz et al., 2017) to recapitulate dynamics in the microfluidic device and account for cell division, input diffusion and cell-to-cell variability; further information for the agent-based simulation settings is reported in the Methods section. Two different closed loop control experiments were simulated: 1) set-point regulation of GFP to 50% (Figure 2A and Video S1A) of its dynamic range, and 2) signal tracking, where the reference value changed from 40% to 80% mid-way through the experiment (Figure 2B and Video S1B). We used ISE (the integral square error, see Methods for further details) as a metric to evaluate the performance of the control strategy; ISE penalises big variations in time of the mismatch providing a reliable assessment of control performance.

**Figure 2.**
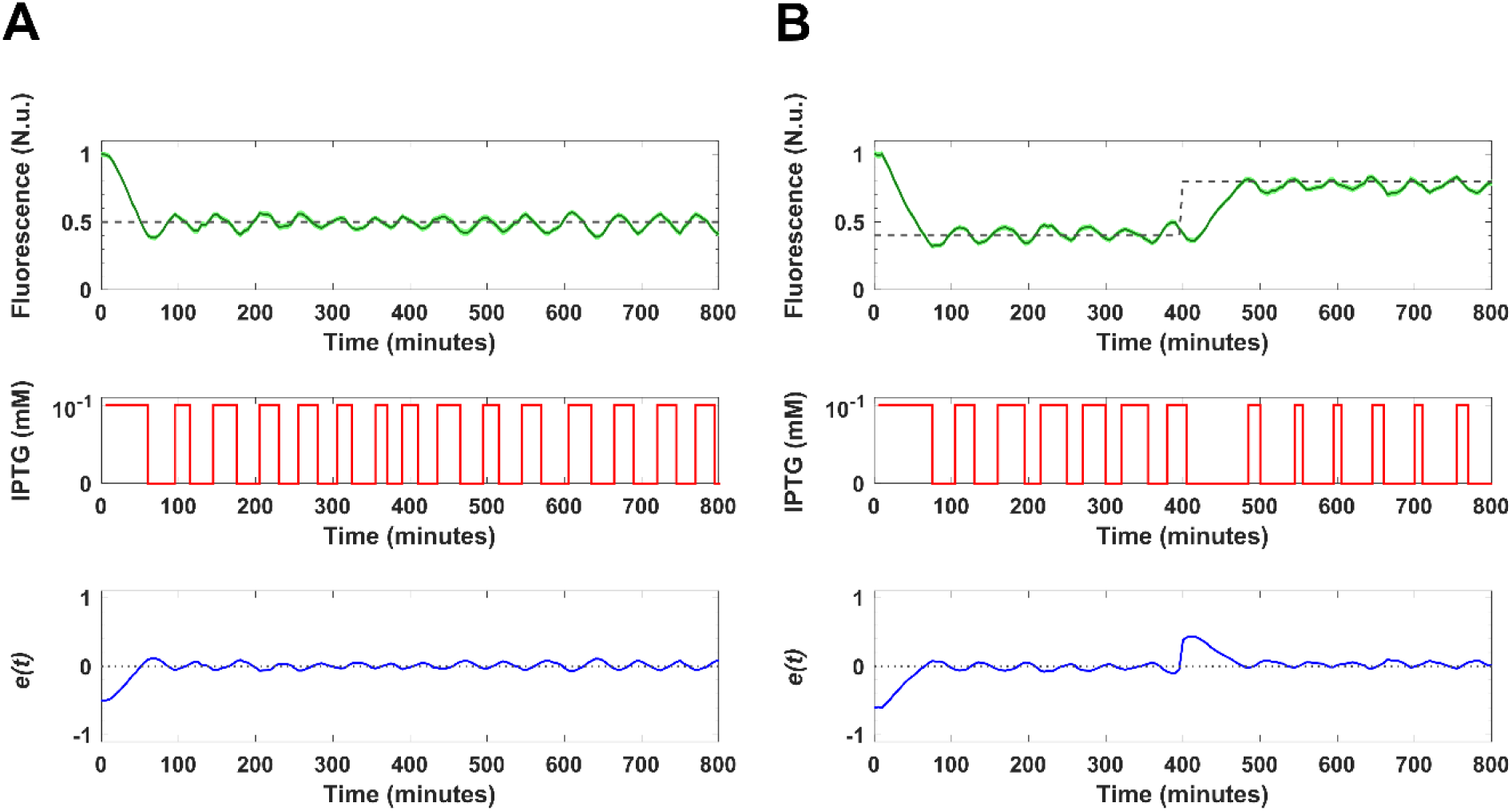
*In silico* set-point and tracking of the molecular titration motif. Simulations were carried out in the agent-based simulator BSim to track GFP expression across the cell population (*y* in green) under a relay control strategy. The simulations were performed with a constant input of 10^−2^ mM AHL. The desired GFP fluorescence was set to **A)** 50% of the dynamic range of fluorescence (*y*_*ref*_ in dashed grey) for the length of the control experiment. **B)** For the first 400 mins of the experiment, the desired GFP fluorescence was set to 40% of the dynamic range of fluorescence, which was then switched to 80% for the remaining 400 mins (*y*_*ref*_ in dashed grey). The input signal (10^−1^ mM IPTG in red), was delivered to the cells when the error (*e*, in blue) between the desired GFP value and the actual GFP was negative. The control input was removed when the calculated error was positive. Control begins at t=0 min. Prior to the *in silico* and *in vivo* experiments start, a calibration phase was performed for the calculation of the minimum and maximum fluorescence values for the tested conditions, see Methods. The integral square error (ISE) was calculated to evaluate the performance of the control strategy, for experiment A, ISE = 7.79 and for experiment B, ISE = 18.15. The standard error of the mean of *y* (SEM) is represented by the narrow shaded region in lighter green.

For both control goals, we observed that the GFP output was driven to the desired level after the first 50 minutes of the experiment and successfully tracked the desired value for the rest of the experiment, with oscillations around the set-point due to the chosen control algorithm (Video S1A and S1B). The *in silico* experiments also show that the ISE for both regulation and tracking experiments remains relatively low (ISE = 7.79 for regulation and ISE = 18.15 for tracking), confirming the effectiveness of the control strategy.

To validate the strategy *in vivo*, we coded the relay controller into the PC running the control experiment. Fluorescence output was sampled every 5 minutes and the control input adjusted accordingly. Specifically, based on the computed mismatch between the sensed and the desired fluorescence values, inducers were either added to, or removed from, the cells by the action of actuators able to automatically change the heights of motorized syringes. The supply and removal of the control input was achieved using an automated IAI Actuation System (see Methods for further details).

Figure 3A (quantification of the experiment in Video S2) and Supplementary Figures S2A andS2D show the effectiveness of the control system for the regulation of fluorescence to a 50% set-point across replica experiments. Due to the initialization phase, the control experiment began with the cells at their highest level of GFP fluorescence. The cell population took approximately 50-100 minutes of 10^−1^ mM IPTG exposure to reach the desired 50% fluorescence set-point. After this point the IPTG control input was modulated by the controller so as to maintain the cells fluorescence around the desired set-point until the end of the experiment at t=800 min. The effectiveness of the control action is confirmed by comparing the ISE of the cells in the controlled chamber with the ISE of cells hosted in two neighbouring chambers that received the same control input but in open loop (Supplementary Table S2).

**Figure 3.**
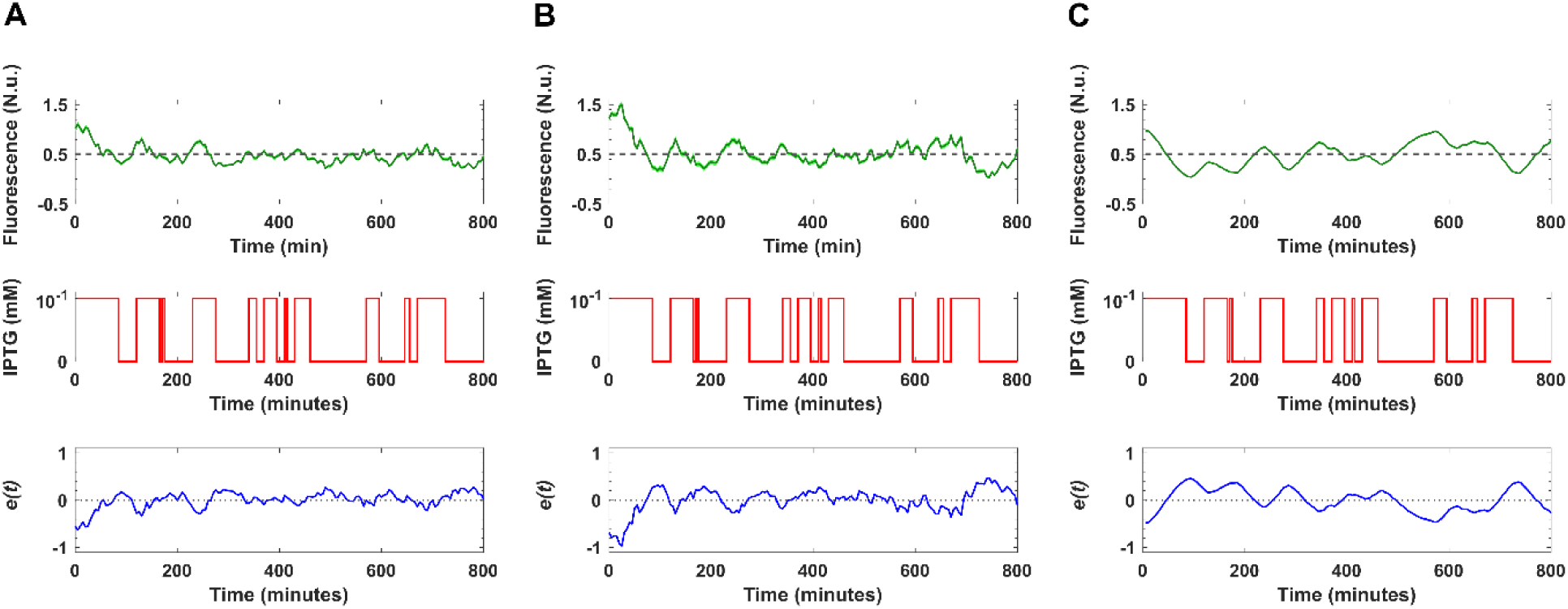
*In vivo* set-point control on the molecular titration motif (50% output). Experiments were performed by considering IPTG modulation with a constant input of 10^−2^ mM AHL. **A)** Real-time control of the GFP fluorescence output from cells inside a single chamber. The desired GFP fluorescence was set to 50% of the dynamic range of fluorescence (*y*_*ref*_ in dashed grey) and is plotted alongside experimentally quantified GFP fluorescence (averaged across the cell population; *y* in green) for the length of the control experiment. The input signal (10^−1^ mM IPTG in red), was delivered to the cells when the error (*e*, in blue) between the desired GFP value and the actual GFP was negative. The control input was removed when the calculated error was positive. **B)** Cells in a neighbouring chamber inside the same microfluidic chip were monitored as an open loop control experiment, since quantification of fluorescence from this cell population did not form part of the feedback control. **C)** *In silico* experiment carried out in the agent-based software platform BSim to track the GFP expression across the cell population (*y* in green) using the same 10^−1^ mM IPTG input recorded during the external control experiment shown in panel A. The simulations were performed with a constant input of 10^−2^ mM AHL. The standard error of the mean of *y* (SEM) is represented by the shaded region. The integral square error (ISE) was calculated for the controlled chamber (22.84), the uncontrolled chamber (51.14) and the *in silico* simulation (45.46). Control begins at t=0 min. Prior to the start of the experiment, a calibration phase was performed for the calculation of the minimum and maximum fluorescence values for the tested conditions, see Methods.

In the set-point experiment shown in Figure 3, the ISE value in the controlled chamber (equal to 22.84) was found to be consistently smaller than that in these other uncontrolled chambers (Figure 3B) where it was up to 2.2-fold higher. The same behaviour was also observed in all replicate experiments, where the ISE values in the uncontrolled chambers (Supplementary Figures S2B and S2E) were found to be even higher being approximately 3- or 4-fold greater than in the controlled chamber (Supplementary Figure S2A, see Supplementary Table S2 for an overview).

A further validation of the experimental observations was obtained by using BSim to simulate the cells’ response in open loop upon provision of the same control inputs recorded during the *in vivo* experiments. Consistent with *in vivo* results, the control error is higher when the system is simulated in open loop (ISE = 45.46, 33.11 and 46.11 in the three replicate experiments; see Figure 3C and Supplementary Figures S2C and S2F) as compared to the aforementioned simulations in closed loop (ISE = 7.79, Figure 2A), confirming the effectiveness of the controller for set-point regulation.

We complemented the regulation experiments described above with a set of tracking experiments where the desired fluorescence set-point (*r*) was initially set to 40% and then increased to 70% or 80% during the experiment at t=400 min. As shown in Figures 4A (recapitulating the experiment shown in Supplementary video S3), the external control input drove GFP fluorescence to 40% of maximum GFP within the first 50-100 minutes of the control experiment. After 400 minutes, the desired fluorescence set-point was switched to 70% or 80%. Cells took approximately 120 mins to reach the new desired set-point in contrast to the 80 minutes obtained from the closed loop BSim simulations, and then tracked the new set-point. The overall ISE was computed to be 30.72 for replicate 1, 69.06 for replicate 2 and 53.68 for replicate 3 (Figure 4A, Supplementary Figures S3A and S3D, respectively).

**Figure 4.**
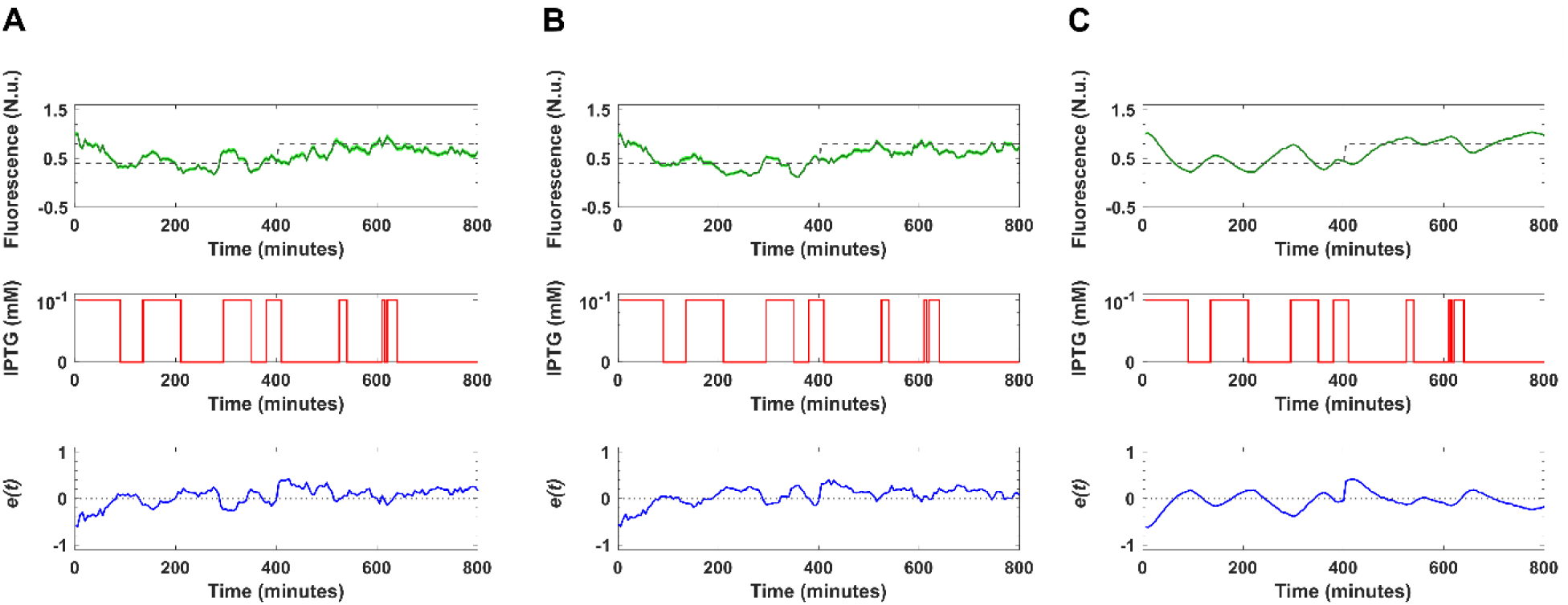
*In vivo* tracking control experiment on the molecular titration motif (40% - 80% output). Experiments were performed with a constant input of 10^−2^ mM AHL. **A)** Real-time control of the GFP fluorescence output from cells inside a single chamber. For the first 400 minutes of the experiment, the desired GFP fluorescence was set to 40% of the dynamic range of fluorescence, which then switched to 80% for the remaining 400 minutes (*y*_*ref*_ in dashed grey). This is plotted alongside experimentally quantified GFP fluorescence (averaged across the cell population; *y* in green) for the length of the control experiment. The input signal (10^−1^ mM IPTG in red), was delivered to the cells when the error (*e*, in blue) between the desired GFP value and the actual GFP was negative. The control input was removed when the calculated error was positive. **B)** Cells in a neighbouring chamber inside the same microfluidic chip were monitored as an open loop control experiment, quantification of fluorescence from this cell population did not form part of the feedback control. **C)** *In silico* experiment carried out by BSim to track the GFP expression across the cell population using the same 10^−1^ mM IPTG input recorded during the external control experiment shown in panel A. The simulations were performed with a constant input of 10^−2^ mM AHL. The standard error of the mean of *y* (SEM) is represented by the shaded region. The integral square error (ISE) was calculated for the controlled chamber (30.72), the uncontrolled chamber (29.87) and the *in silico* simulation (26.69). Control begins at t*=*0. Prior to experiment start, a calibration phase was performed for the calculation of the minimum and maximum fluorescence values for the tested conditions, see Methods.

As before, ISE values obtained for the controlled chambers were compared with those computed for two other neighbouring chambers that were receiving the same control input in open loop. The ISE values obtained in the controlled chambers were lower than those in the uncontrolled chambers in two of the three replicate experiments (see Figures 4A and B, Supplementary Table S2). Specifically, in two of the three replicate experiments, ISE values in the uncontrolled chambers were observed to be approximately 1.5-fold and 5-fold higher than in the controlled chamber and in the third replicate the ISE values were comparable in the controlled and uncontrolled chambers, see Supplementary Figures S3A, S3B, S3D and S3E, and Supplementary Table S2.

Feedback control was in general also observed to increase cell coherence as shown by computing the standard deviation of the error signal across the cell population, which in most cases is smaller in the controlled chambers than in the uncontrolled ones, see Supplementary Table S2.

## Discussion

We have shown that it is possible to modulate in real-time the response of a testbed antithetic motif by means of external feedback control in microfluidics. Closed loop control using a set-point reference signal was effective both *in silico* and *in vivo* and the system also responded well to a dynamic reference signal *in silico.* Experiments tracking a dynamic reference signal *in vivo* produced inconsistent results, suggesting that there may be elements associated with transient disruptions to biological systems that are not currently captured by our model, and which might require more sophisticated control strategies.

The module is based on an orthogonal *σ* /anti-*σ* pair that was firstly presented in our previous work (Annunziata et al., 2017) and is similar to those recently used for the implementation of antithetic feedback controllers *in vivo* and *in vitro* (Aoki et al., 2019, Agrawal et al., 2019). The experimental validation we provide here integrates the quasi-static experimental investigation of the *σ* /anti-*σ* motif of interest which was started in (Annunziata et al., 2017) complementing it with microfluidics data that gives useful information on typical response times and the overall behaviour of the module when stimulated in real-time.

Its characterization in microfluidics will be invaluable for future implementation of *in vivo* control strategies such as antithetic integral feedback control. Another potential application is for the realization of the *in vivo* multicellular control we proposed in earlier work (Fiore et al., 2017) where the module we studied in this paper is the core of a controller population taming the behaviour of some target cell population of interest. In that scenario, IPTG is used to provide a, possibly time-varying, reference signal to the controller cells that need to respond in real-time by comparing its level with that of an AHL signal acting as a proxy of the output of the target population. The results reported in this paper show a response time of about 100 minutes which is compatible with the time-scale of the multicellular control experiments described in (Fiore et al., 2017) indicating that the *σ* /anti-*σ* motif can indeed be used in that context.

From a control viewpoint, our results provide further confirmation that real-time external control of bacterial cell populations can be achieved *in vivo* by appropriately varying inducer molecular input to the cells hosted in a microfluidics platform. In this respect, our results complement those in (Lugagne et al., 2017) which, to the best of our knowledge, is the only other work in the literature achieving *in vivo* control of bacterial populations via microfluidics.

During the analysis of our *in vivo* experimental data, we encountered a considerable degree of cell-to-cell variability with respect to GFP production. We determined that this observation was not an artefact of our segmentation algorithm by analysing the same images with a separate deep learning segmentation algorithm (data included in a submitted manuscript). We concluded the source of this variance arose from inherent stochasticity found within biological systems (Spudich and Koshland, 1976, Rao et al., 2002). Several studies have identified sources and mechanisms behind biological noise and it is widely accepted that factors such as gene copy number, availability of transcription factors and cellular machinery, cell stress, and cell life cycle can all result in phenotypic variability despite identical genomes (Arkin et al., 1998, Becskei et al., 2005, Cai et al., 2006, Raj and van Oudenaarden, 2008, Takhaveev and Heinemann, 2018). Specifically, studies using *E. coli* have explicitly noted phenotypic heterogeneity within clonal populations under homogenous conditions (Nikolic et al., 2017, Nikolic et al., 2013, Keegstra et al., 2017). Our system involves the coordination of gene expression over three different plasmids, each with medium/low copy number, resulting in multiple aspects where small component changes have the potential to lead to large variance in expression (Jahn et al., 2016). Despite cell-to-cell variability, we were still able to accurately regulate the mean fluorescence of the population using a simple algorithm to direct the behaviour of our antithetic motif. The large amount of cell-to-cell variability can explain the better response observed in regulation experiments where the set-point is kept constant all through the experiments than in tracking experiments where it is changed halfway through and therefore triggers a new transient response from the cells.

Future work will be aimed at characterizing scale effects in the response of the module when experiments are carried out in larger vessels such as the turbidostat we are currently designing (Guarino et al., 2019). Also, ongoing work is using the module we characterized to close *in vivo* the feedback loop across two bacterial populations which was designed and tested *in silico* in our previous work (Fiore et al., 2017).

## Acknowledgments

The authors wish to acknowledge the invaluable help of Dr Ryan Johnson and Prof Jeff Hasty from UC San Diego for providing the design of the microfluidics chip used to carry out the experimental work described in this paper. The authors thank Dr Dan Rocca and Dr Elisa Pedone for support with the microfluidics platform set-up, and Dr Mark Jepson and Alan Leard (Wolfson Imaging Facility, University of Bristol) for supporting live-cell imaging experiments.

This work was supported by BrisSynBio, a BBSRC/EPSRC Synthetic Biology Research Centre (BB/L01386X/1) to CG, NS, LM and MdB, by EC funding H2020 FET OPEN 766840-COSY-BIO to LM and MdB, and by EPSRC funding to LM (EP/R041695/1 and EP/S01876X/1).

## Author Contributions

BS optimized the microfluidics platform and carried out the *in vivo* experiments; CGZ performed the modelling and *in silico* experiments and contributed to the segmentation algorithm; LP developed the segmentation algorithm and supported the experimental investigation; DS performed the agent-based simulations in BSim; all authors analysed and discussed the data; MdiB, NS, CG and LM conceived the study; BS, CGZ,LM, NS and MdiB wrote the manuscript.

## STAR Methods

### Experimental model and subject details

#### Strains and constructs and media

The *E. coli* strain and plasmid constructs used in this report have been previously described in (Annunziata et al., 2017). The reference-comparator system was hosted within *E. coli* MG1655 (*λ-, rph-1*) (Guyer, 1981) and was composed of three plasmids, a sigma producing plasmid (pLuSb, chloramphenicol resistant), an anti-sigma producing plasmid (pVRa_LacASb_Flag, ampicillin resistant) and a GFP reporter plasmid (pVRb_ssrA, kanamycin resistant). The pLuSb plasmid encoded the LuxR gene, controlled by the lacI promoter, downstream, the sigma molecule was placed under the control of the plux promoter (AHL responsive). The pVRa_LacASb_Flag plasmid encoded the LacI gene, controlled by the lacI promoter, downstream, the anti-sigma molecule was controlled by lacUV5 promoter (IPTG responsive). The pVRb_ssrA plasmid encoded the sfGFPssrA gene, controlled by the p20_992 promoter, whose activity was regulated by the sigma/anti-sigma pair. To ensure fast dynamics of expression and degradation the sigma, anti-sigma and GFP proteins contained a ssrA tag (AANDENYALAA). The pLuSb and pVRa_LacASb_Flag are medium copy number plasmids and pVRb_ssrA is a low copy number plasmid.

All cell culture incubations and microfluidics experiments were performed using Luria-Bertani (LB) medium (MP Biomedicals) supplemented with kanamycin (50 µg/mL), ampicillin (100 µg/mL) and chloramphenicol (25 µg/mL). All antibiotics were supplied by Sigma-Aldrich.

AHL (N-(B-Ketocaproyl)-L-Homoserine Lactone from Sigma-Aldrich, cat # K3007) was dissolved in water, filter-sterilized and added to the LB growth medium at the indicated concentrations. IPTG (Isopropyl β-D-1-thiogalactopyranoside, supplied by Sigma-Aldrich, cat # I5502) was dissolved in water, filter-sterilized and added to LB medium at the indicated concentrations.

## Method details

### Microfluidic device and fabrication

The microfluidic device design used in this report was developed by Mondragón-Palomino and colleagues at the University of California, San Diego (Mondragon-Palomino et al., 2011). A replica of the silicon mould (containing eight identical designs) was donated to our group. Soft lithography was used to form the microfluidics devices using methods described in Ferry et al, 2011 (Ferry et al., 2011), which were adapted to meet our specific requirements. First a 10:1 ratio of silicone elastomer base and curing agent (SYLGARD 184, Ellsworth Adhesives Limited) was weighed into a large plastic weigh boat and mixed vigorously for 3 minutes. A foil tray was built around the silicon mould and the mixture was poured onto the patterned side of the mould and placed inside a degassing chamber until all air bubbles were removed from the mixture. The PDMS mixture was cured onto the silicon mould by baking at 80°C for 2 hours and then allowing to cool. The PDMS layer was carefully peeled away from the silicon mould and the individual devices cut out using a scalpel. A biopsy puncher (0.75 mm, Darwin Microfluidics) was used to punch holes through the device to create the ports. The PDMS was washed with isopropanol followed by double distilled water (ddH_2_O) and dried using an air gun. Scotch tape was rubbed onto the surface of the device and peeled away 5 consecutive times to remove any remaining debris. A thin glass slide (150 µM, Hirschmann Deckgläser) was prepared by cleaning with acetone, methanol, isopropanol and ddH_2_O and then dried using an air gun. Finally, the patterned surface of the PDMS and the cleaned glass slide were exposed to plasma using a Plasma Asher machine (Diener Zepto) set to 50 % power, for 30 seconds and then brought into contact. The bonded devices were incubated at 80°C overnight to strengthen the bond.

### Preparation of the microfluidic device

For a schematic representation of the microfluidic device please refer to Mondragón-Palomino et al 2011 (Mondragon-Palomino et al., 2011). Before the experiment start, a wetting protocol was performed on the microfluidic device to remove any air bubbles and debris from inside the PDMS. Using tape, the microfluidic device was gently secured on top of a thick glass slide. Approximately 50 cm of Tygon laboratory tubing (ND 100-80, Saint Gobain) was cut. A 2.5 mL syringe was filled with approximately 1 mL of fresh media (and 0.075% Tween-20, Sigma-Aldrich) and attached to one end of the tubing via a 23 gauge needle (Microlance), the other end was attached to the ‘R’ port of the microfluidic device via a 23 gauge bent metal pin (Metcal). Manual pressure was applied using the syringe plunger to push the media through the device. Once media had pushed up through ports B, I and W2, the bent pin was transferred into the W2 port and the manual pushing continued until media pushed through the W1 and C ports. The device was then inspected under the microscope for any remaining air bubbles and debris using the 40X objective (see below for microscope details). If air bubbles and debris remained, the wetting protocol was repeated. This inspection was also used to check the integrity of the device, making sure that all features had formed correctly. The wet microfluidic device was placed on top of the microscope incubator box (approximately 25 cm above stage height), ready for connection to fluidic lines.

### Culture preparation

One colony of the reference-comparator strain was selected to seed 5 mL of LB media with antibiotics and grown overnight (approximately 16 hours) at 37 °C with shaking (200 rpm). 300 µL of the overnight culture was used to seed 300 mL of fresh LB medium (plus antibiotics). This culture was grown to an optical density of 600 nm of 0.3 (OD_600nm_). The culture was then split into 6 × 50 mL aliquots inside 50 ml Falcon tubes, centrifuged at (2,200 *x g*) for 15 minutes and resuspended in 1.5 mL of fresh LB medium supplemented with 0.075 % Tween-20 (Sigma-Aldrich) and antibiotics.

### Preparation of lines

Six lengths of Tygon tubing were cut (3 × 100 cm; 3 × 200 cm, ND 100-80, Saint Gobain). For each length of tubing, a 23 gauge Luer needle was attached to one end. A 23 gauge bent metal pin was attached to the other end. Luer needles were connected to the Luer slip of a 10 ml syringe (plunger removed). Before bringing the Luer needle and syringe into contact, the head of the Luer needle was filled with approximately 200 µL of the appropriate media to prevent air bubbles forming inside the connection. After connecting, the Luer syringe was filled with 4 mL of appropriate media (detailed below). The syringe was then gently flicked to remove any remaining air bubbles. The connected line then filled with media by gravity flow. Once media began to exit the line through the bent pin, a small amount of tape was used to attach the pin to the side of the syringe to prevent any more flow whilst the other lines were prepared.

Lines cut to 200 cm lengths were used to connect the B, R and I ports. Lines cut to 100 cm lengths were used to connect the W1, W2 and C ports. The 10 mL syringes were filled with the appropriate media and connected to microfluidic device (on top of the microscope box) in the order described below. The height above the stage which the syringes were placed is noted inside the parentheses:

**W2:** 4 mL of fresh media plus 0.075% Tween-20 and antibiotics (30 cm)

**W1:** 4 mL of fresh media plus 0.075% Tween-20 and antibiotics (37 cm)

**B:** 4 mL of fresh media plus 0.075% Tween-20, antibiotics, 10^−2^ mM AHL and 0.4 µL of Sulforhodamine B dye (60 cm)

**R:** 4 mL of fresh media plus 0.075% Tween-20 and antibiotics (60 cm)

**I:** 4 mL of fresh media plus 0.075% Tween-20, antibiotics, 10^−2^ mM AHL and 10^−1^ mM IPTG (60 cm)

**C:** 1.5 mL of cell culture (32.5 cm)

### Loading cells into the device

Once all lines were attached, the microfluidic device was transferred down to the microscope stage and gently secured with tape. Using phase contrast live imaging under the 40X objective, cells were loaded into the trapping chambers of the device via the C port. If cells didn’t immediately flow from the C port, the line was gently flicked and the heights of the W1 and W2 syringes were lowered slightly to encourage flow. Once a steady flow of cells into the device was established, the heights of the W1 and W2 syringes were increased slightly to direct the flow of cells towards the trapping region. The line connected to the C port was then gently flicked to encourage cells to enter and become trapped inside the chambers of the microfluidic device. The device was determined as sufficiently loaded when approximately 10 or more chambers contained cells. After this point, the W1, W2 and C ports were lowered to a height of 14 cm above stage height. To balance to ports and prevent clogging, 1 mL of culture was removed from the C port (whilst the line was still connected to the device) and replaced with 3.5 mL of fresh media (0.075% Tween-20 plus antibiotics). Syringes connecting the B and I ports were then attached to the automatic actuators and raised to a height of 195 cm and 233 cm respectively. The R port was kept at a height of 60 cm.

### Phase contrast and fluorescence microscopy

For each experiment, time-lapse fluorescence microscopy was performed using an inverted widefield fluorescence microscope (Leica) fitted with an Andor iXon Ultra digital camera. The microfluidic device was secured onto the stage of the microscope and enclosed inside an incubation chamber set to 37°C (Pecon). The microscope was programmed to take images of cells growing inside three different trapping chambers every 5 minutes. At each time point a phase contrast image (PhC), a green fluorescence image and a red fluorescence image were acquired for each trapping chamber. Green fluorescence images were used for the detection of GFP and red fluorescence images were used for the detection of the Sulforhodamine B dye. All images were acquired using the 100 X objective. To prevent any problems of drift, the Leica autofocus setting was utilised. Exposure times were set to 100 ms for the PhC and green spectra and 150 ms for the red spectra.

### Segmentation for bacterial cells

The algorithm to segment the microscopy images aims at distinguishing the foreground (bacterial cells) from the background in each image of the time lapse experiment. The raw images were pre-processed as follows before running the segmentation routine. Specifically during the pre-processing stage: 1) images were cropped to the area where individual cells had to be segmented; 2) an interpolation was performed to set the proper size of the image and 3) the cropped image was filtered to minimize noise and blurriness. The filters used to pre-process the image were implemented by the MATLAB routines ‘wiener2’ (2-D adaptive noise-removal filtering) and ‘medfilt2’ (2-D median filtering). The neighbourhood sizes used to implement those filters were set to 9 and 3, respectively. Furthermore, we used the MATLAB image processing function ‘adapthisteq’ to enhance the contrast of the grayscale image.

The algorithm then applied Otsu thresholding (Vala and Baxi, 2013) to binarize the grey scale phase contrast image and used morphological operators such as erosion and dilation (MATLAB routines ‘imerode’ and ‘imdilate’) to calculate the global area where cells are allocated, and found cells centres and edges (local thresholding). Local thresholding was applied to find cell centres and edges to segment individual objects (single cells). All the functions and filters were taken from the MATLAB image processing toolbox.

To calculate the final mask a refinement stage was carried out to delete any small object which was not identified within the range of the cell morphology. This range was calculated by choosing the average area from all the objects and excluding the objects smaller than the 50% of that value. Finally, the algorithm identified the bigger objects and separated them into individual cells. For doing this, the algorithm chose the objects, which have an area twice the average area, calculated over all the objects present in the mask. These larger objects were further segmented to identify individual cells.

The final mask contained the boundaries and interiors of every segmented cell; this mask was overlapped over the green field image to calculate the fluorescence corresponding to the pixels in the interior of the segmented cells. To calculate the fluorescence at every sampling time, the background value was subtracted from the fluorescence quantification. The background was calculated as the mean of the pixels acquired from cropping an area of the image where there were no cells, which was in the main channel of the device, within the green field image. A threshold value was set to include only cells whose fluorescence was within a maximum range and to discard fluorescence from outliers cells. The threshold was calculated as the fluorescence value of the cell exhibiting the highest fluorescence in the set of 60 images acquired during the 300 min initialization phase preceding each control experiment. The final average fluorescence value was then computed as the mean of the fluorescence exhibited by all the objects left in the final mask after removing those above the threshold.

### Actuation

An IAI Actuation System (LC Automation) controlled by Dial-A-Wave (DAW) software (Ferry et al., 2011) was used for the automatic supply and removal of chemical signals for both open and closed loop experiments. Syringes connected to the B and I ports of the microfluidic device were attached to vertical actuators mounted onto a wall. For closed loop relay control, our MATLAB algorithm controlled the heights of the actuators via the DAW software by continually re-writing text files, which the DAW software referenced for height positioning.

### Parameterisation of the model

Our existing mathematical model previously developed for batch culture simulations (Annunziata et al., 2017) was parameterised to reflect the conditions of our microfluidics experiments. This was done using deterministic simulations using ODE45 solver from MATLAB to fit the model predictions to the experimental data collected from *in vivo* open loop microfluidics experiments. Firstly, *in vivo* data was collected during open loop experiments with a constant input of 10^−2^ mM AHL, and 10^−1^ mM IPTG supplied to or removed from the cells every 4 hours. Next, a search was performed to find the parameters that gave the model outputs best matching the experimental observations. The genetic algorithm function (ga) from the MATLAB optimisation toolbox was used find the optimal value of the following objective function, which calculates the mean square error (MSE) between the model and experimental data points:

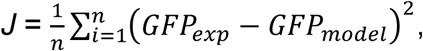

where *i* is the current sampling time and *n* is the number of samples used for the fitting, *GFP*_*exp*_ is the normalized fluorescence measured *in vivo* and *GFP*_*model*_ is the normalized fluorescence predicted by the *in silico* model (Supplementary Figure S1).

In Supplementary Figure S1, we show that the model with the new optimized parameters reported in Supplementary Table S1 can capture the fluorescence values detected during *in vivo* experiments. To further validate the predictive ability of our model, we used additional *in vivo* open loop experimental data acquired using different concentrations of signalling molecules. In particular, Supplementary Figure S1B shows the comparison between model and experiments when the control input 1 mM AHL was supplied to or removed from the cells every 3 hours, in the absence of IPTG. Additionally, Figure S1C shows the model predictions against experimental results when a control input changed from 10^−2^ mM AHL with no IPTG to 10^−4^ mM AHL with 10^−1^ mM IPTG every 4 hours.

### Control strategy

*In silico* and *in vivo* control experiments were designed to keep GFP fluorescence output from the cells at a fixed set-point or track a time-varying reference. AHL at a concentration of 10^−2^ mM was continually supplied to the cells to maintain production of the sigma factor, while IPTG was used as a control input by switching between 0 mM and 10^−1^ mM. The control experiment began at *t* = 0 and lasted 800 minutes. Prior to the control start, a calibration phase was used to calculate the minimum and maximum GFP. To calculate the minimum GFP value, cells were exposed to 10^−1^ mM IPTG for 300 minutes, GFP values obtained in the last 30 minutes of this phase were averaged and normalised to 0. To calculate maximum GFP, IPTG was removed for 300 minutes. GFP values obtained in the last 30 minutes of this phase were averaged and normalised to 1. After calibration, relay control was used to modulate the supply and removal of the control input, in order to regulate the GFP to the desired output. This form of control requires only the calculation of the control error (*e*) between the desired reference value (*r*) and the actual GFP output (*y*) at each time point (Åström and Murray, 2010). This control strategy was first simulated *in silico* before being implemented *in vivo*.

### In silico closed loop control experiments

We simulated the control experiments by stochastic simulations using BSim, an agent-based simulator able to mimic realistically microfluidic experiments (Matyjaszkiewicz et al., 2017). Specifically, this simulator considers cell reproduction, spatial distribution, cell geometry and movements, the spatial distribution of the external molecules, the diffusion and degradations of the chemicals and the spatial applications of the control inputs. In our simulations, we mimic the growth of the monolayer *E. coli* population (approximate 200 cells) inside in a chamber with the same characteristics of the one we used for *in-vivo* experiments.

When a cell divides, we modify the values of production rates of protein species (*σ, σ*_*α*_, *σ:σ*_*α*_ and *GFP*) to take into account asymmetric plasmid copy numbers in the daughter cells. These parameters were extracted from a uniform distribution in the interval (*n*±25%), where *n* is the nominal value of the maximal rate of production for (*χ*_1,*i*_), *i= σ, σ*_*α*_, *σ:σ*_*α*_ and *GFP* (Supplementary Table S1). This allowed us to reproduce population-level heterogeneity observed experimentally.

We used the pseudo reactions described in the deterministic model (Equations 1-4, main text) to simulate our system stochastically. The propensities are taken from the production and degradation terms of a set of stochastic differential equations (SDEs). The SDE system was solved numerically using the Euler Murayama algorithm. All the simulations were performed with a laptop processor Intel Core i7-8850H CPU@2.6Hz.

### In vivo closed loop control experiments

*In vivo* control experiments were realised using the experimental set up described above. The control input of 10^−1^ mM IPTG was supplied via the syringe attached to the ‘I’ port of the microfluidic device. Both the syringes attached to the ‘B’ and ‘I’ ports contained 10^−2^ mM AHL. Time lapse microscopy was used to capture the phase contrast and fluorescence images as described above every 5 minutes. The MATLAB controller performed the online bacterial segmentation algorithm in order to quantify the GFP fluorescence from the cells and calculate the control error. Based on this control error the IAI was instructed to either swap the heights of the B and I syringes or keep them the same.

## Quantification and statistical analysis

### Statistical moments

The mean and standard deviation of the fluorescence signal were calculated from the data obtained via the segmentation algorithm described earlier. All computations were carried out using MATLAB.

### Error signal analysis

To compare the value of the error among all experiments. The mean square error (MSE) of the error signal *e(t)* was calculated for each experiment as follows:

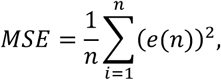

where *n* is the number of sampling times over the control experiment.

### Performance evaluation

We chose the Integral square error (ISE) to evaluate the performance of the relay controller *in vivo* and *in silico* experiments. The ISE is calculated over every signal error *e(t)* as the integral over time of the control error squared. Specifically,

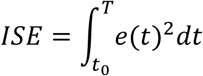

where *t*_*0*_ is the initial time and *T* is the duration of the experiment.

## Data and code availability

All the raw data can be downloaded from: https://github.com/diBernardoGroup/InVivoFeedbackControl

The segmentation code is available upon request via email from

