## Supplementary Information for "*In vivo* Feedback Control of an Antithetic Molecular-Titration Motif in *Escherichia coli* using Microfluidics"

### Supplementary information – Submission Manuscript External Control

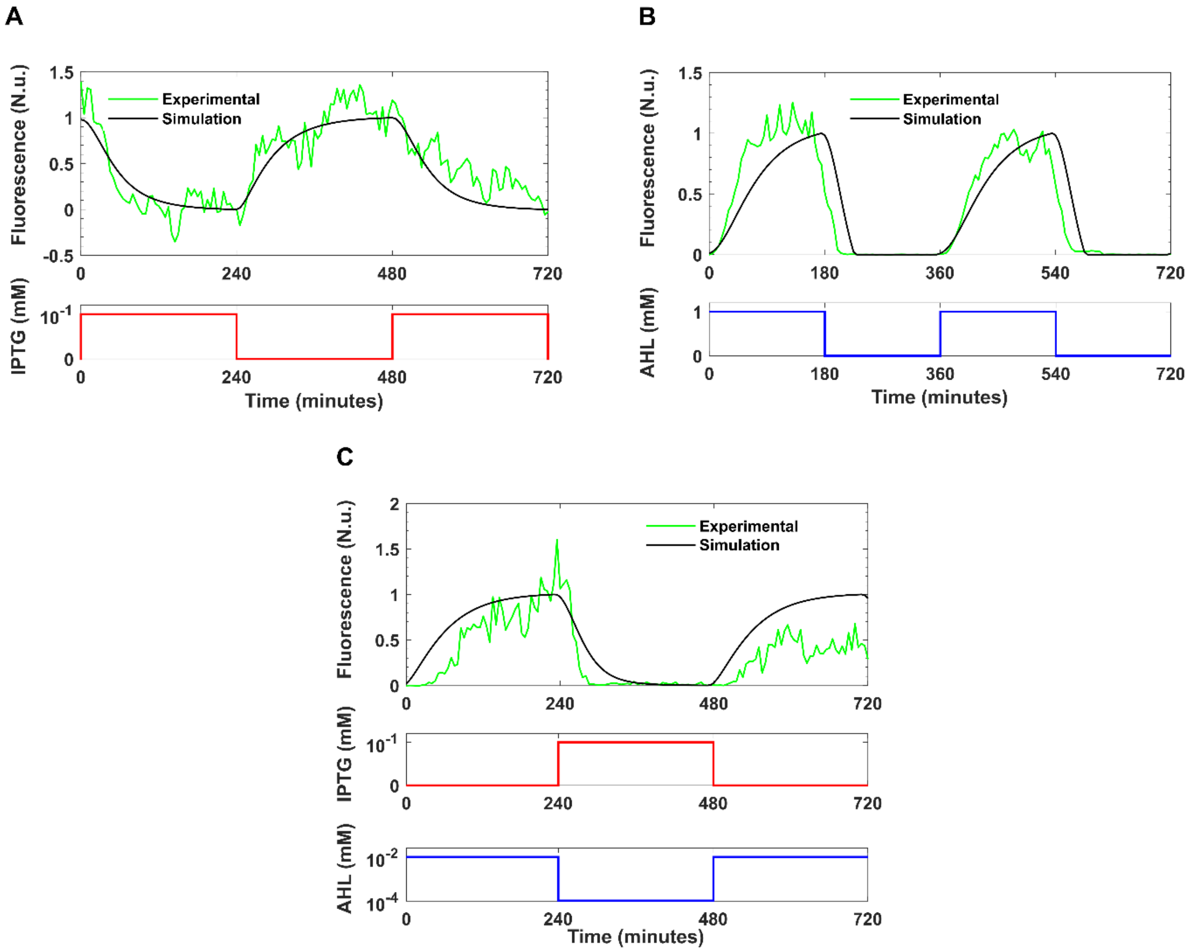


**Figure S1. Fitting and validation of the model.** GFP expression over time measured in *in vivo* (green) and *in silico* (black line) open loop experiments. **A)** For the sake of fitting, the input is selected as a square wave of IPTG (red) changing from 0 to $10^{-1}$ mM every 4hrs with constant AHL set at $10^{-2}$ mM. For the sake of validation: **B)** The input is a square wave of AHL (blue) changing from 0 to 1 mM every 3hrs and **C)** The inputs are a square wave of IPTG (red line) switching from 0 to $10^{-1}$ mM and a square wave of AHL (blue) switching from $10^{-2}$ to $10^{-4}$ mM every 4hrs.

| Parameter | Description | Baseline value (Annunziata et al., 2017) | Optimised value |
|---|---|---|---|
| $\chi_{0,i}$ | Basal rate of production for proteins species $i$ | 54 molecules $min^{-1}$ | |
| $\chi_{1,i}$ | Maximal rate of production for proteins species $i$ | 1080 molecules $min^{-1}$ | |
| $\chi_{0,GFP}$* | Basal rate of production for GFP | | 53.89 molecules $min^{-1}$ |
| $\chi_{1,GFP}$ * | Maximal rate of production for GFP | | 998.57 molecules $min^{-1}$ |
| $K_\sigma$ | Microscopic dissociation constant for σ-regulated promoter | 1.98x$10^4$ molecules per cell | |
| $K_I$ | Microscopic dissociation constant for IPTG-regulated promoter | 90 $\mu$M | 35 $\mu$M |
| $K_A$ | Microscopic dissociation constant for AHL-regulated promoter | 10 nM | |
| $n_\sigma$ | Hill coefficient for σ-regulated promoter | 1.93 | 0.46 |
| $n_A$ | Hill coefficient for AHL-regulated promoter | 0.31 | 0.35 |
| $n_I$ | Hill coefficient for IPTG-regulated promoter | 0.46 | 1.99 |
| $\gamma_{P_i}$ | Rate of translation of protein species i from its mRNA | 0.0277 $min^{-1}$ | |
| $\gamma_D$ | Maximal rate of degradation through ssrA tags | 1080 molecules $min^{-1}$ | 600.2 molecules $min^{-1}$ |
| $c_e$ | Half-maximal concentration for kinetics of ssrA tag based degradation. | 0.01 molecules per cell[1] | 0.08 molecules per cell |
| $k^{+}_{\sigma:\sigma_\alpha}$ | Rate of complex formation (association of σ and $\sigma_\alpha$). | 0.018 molecules $min^{-1}$ | |
| $k^{-}_{\sigma:\sigma_\alpha}$ | Rate of complex dissociation ($\sigma:\sigma_\alpha$ into its constituents) | 0.00018 $min^{-1}$ | |

**Table S1.** Kinetic parameters of the ODE model and their meaning, along with previous values published in (Annunziata et al., 2017) and the final optimised value after an identification procedure was carried out on experimental data described in the main text (See Methods).

*These parameters were optimised according to the identification procedure described in the Methods (Parameterisation of the model).

[1] Note that the model captures aggregate cell behaviour and therefore fractional values are admissible here.

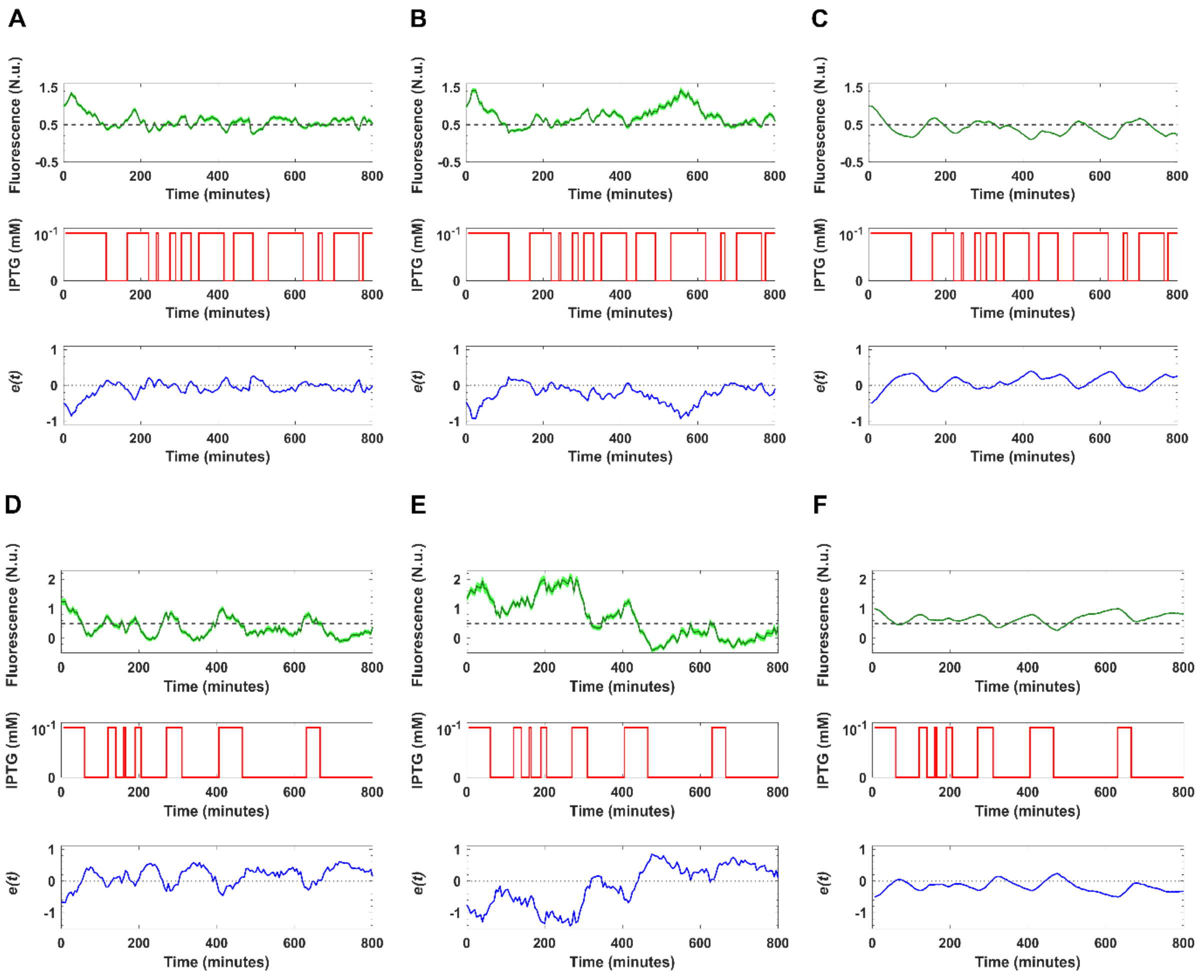


**Figure S2**. **Repeats of *in vivo* set-point control experiment on the molecular titration motif (50% set-point).** Experiments were performed with a constant input of 10$^{-2}$ mM AHL. **A)** Controlled chamber for replicate 2. The desired GFP fluorescence was set to 50% of the normalized fluorescence computed during the initialization stage ($y_{ref}$ in dashed grey). The input signal (10$^{-1}$ mM IPTG in red), was delivered to the cells when the error ($e$, in blue) between the desired GFP value and the actual GFP was negative. The control input was removed when the calculated error was positive. **B)** Uncontrolled chamber for replicate 2 with control input from A. **C)** *In silico* simulation using the same IPTG input recorded during the external control experiment for replicate 2. **D)** Controlled chamber for replicate 3. **E)** Uncontrolled chamber for replicate 3 with control input from D. **F)** *In silico* simulation using the same 10$^{-1}$ mM IPTG input recorded during experiment for replicate 3. The standard error of the mean of $y$ (SEM) is represented by the shaded region. For replicate 2 and 3, the ISE values were calculated for the controlled, uncontrolled and *in silico* simulations **A**= 36.50, **B**= 99.85, **C**= 33.11, **D**= 86.38, **E**= 364.39 and **F**= 46.11 respectively. Control begins at $t=0$. Prior to experiment start, a calibration phase was performed for the calculation of the minimum and maximum fluorescence values for the tested conditions, see Methods.

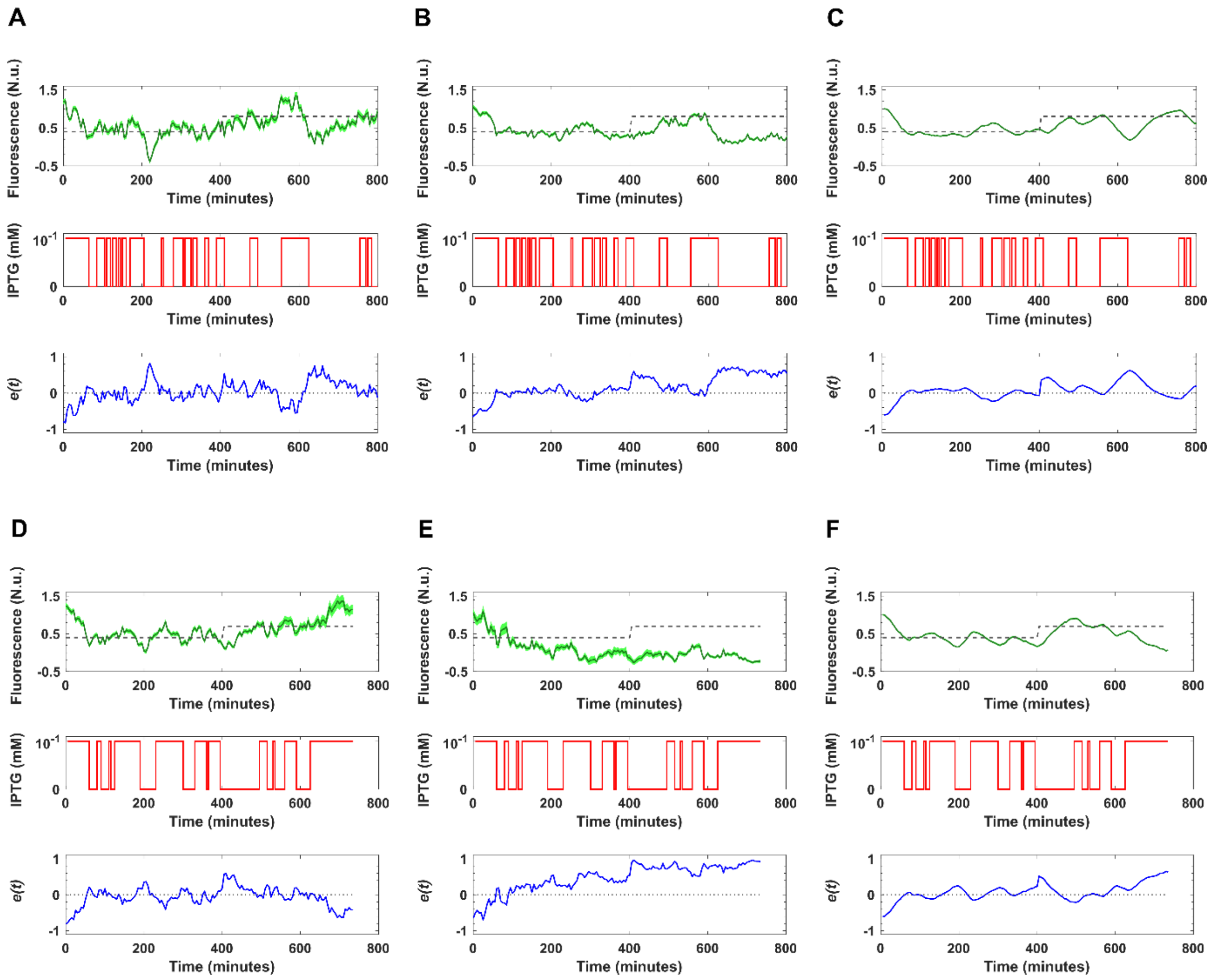


**Figure S3**. **Repeats of *in vivo* tracking control experiment on the molecular titration motif.** Experiments were performed with a constant input of $10^{-2}$ mM AHL. **A)** Controlled chamber for replicate 2, for the first 400 minutes of the experiment, the desired GFP fluorescence was set to 40% of the dynamic range of fluorescence, which then switched to 80% for the remaining 400 minutes ($y_{ref}$ in dashed grey). The input signal ($10^{-1}$ mM IPTG in red), was delivered to the cells when the error (*e*, in blue) between the desired GFP value and the actual GFP was negative. The control input was removed when the calculated error was positive. **B)** Uncontrolled chamber for replicate 2 with control input from A. **C)** *In silico* simulation using the same $10^{-1}$ mM IPTG input recorded during the external control experiment for replicate 2. **D)** Controlled chamber for replicate 3, for the first 400 minutes of the experiment, the desired GFP fluorescence was set to 40% of the dynamic range of fluorescence, which then switched to 70% for the remaining 335 minutes ($y_{ref}$ in dashed grey). **E)** Uncontrolled chamber for replicate 3 with control input from D. **F)** *In silico* simulation using the same IPTG input recorded during experiment for replicate 3. The standard error of the mean of *y* (SEM) is represented by the shaded region. For replicate 2 and 3, the ISE values were calculated for the controlled, uncontrolled and *in silico* simulations **A**= 69.06, **B**= 100.56, **C**= 39.81, **D**= 53.68, **E**= 265.3 and **F**= 43.55, respectively. Control begins at *t=0*. Prior to the control experiment, a calibration phase was performed for the calculation of the minimum and maximum fluorescence values for the tested conditions, see Methods.

| | | Controlled chamber | Uncontrolled chamber 1 | Uncontrolled chamber 2 |
|---|---|---|---|---|
| **Set-point experiments** | **MSE** | | | |
| | Replica 1 | 0.0293 | 0.2121 | 0.065 |
| | Replica 2 | 0.0461 | 0.1247 | 1.6913 |
| | Replica 3 | 0.1088 | 0.4544 | 3.0695 |
| | **Standard Deviation** | | | |
| | Replica 1 | 0.1717 | 0.4492 | 0.2558 |
| | Replica 2 | 0.1958 | 0.2604 | 0.3856 |
| | Replica 3 | 0.2933 | 0.6552 | 1.2734 |
| | **ISE** | | | |
| | Replica 1 | 22.84 | 168.43 | 51.14 |
| | Replica 2 | 36.4985 | 99.8511 | 1355.6 |
| | Replica 3 | 86.3762 | 364.3872 | 2312.4 |
| **Tracking experiments** | | **Controlled chamber** | **Uncontrolled chamber 1** | **Uncontrolled chamber 2** |
| | **MSE** | | | |
| | Replica 1 | 0.0393 | 0.0381 | 0.0567 |
| | Replica 2 | 0.0878 | 0.1272 | 0.1316 |
| | Replica 3 | 0.0754 | 1.0709 | 0.3629 |
| | **Standard Deviation** | | | |
| | Replica 1 | 0.1947 | 0.186 | 0.2389 |
| | Replica 2 | 0.2950 | 0.3131 | 0.3368 |
| | Replica 3 | 0.2689 | 0.6523 | 0.3807 |
| | **ISE** | | | |
| | Replica 1 | 30.72 | 29.8682 | 44.509 |
| | Replica 2 | 69.06 | 100.5634 | 103.4325 |
| | Replica 3[2] | 53.68 | 778.36 | 265.30 |

**Table S2. Analysis of error signals of set-point and tracking experiments *in vivo*.** Mean square error (MSE), standard deviation (SD) and Integral Square Error (ISE) values in real-time closed loop *in vivo* experiments for controlled and uncontrolled chambers. See Methods for MSE, SD and ISE quantifications. In the figures, we chose to display data from the best performing uncontrolled chambers.

[2] Note that this replicate experiment lasted 735 min so the ISE is computed over 735 mins rather than 800 mins. Again, we see that ISE values in the uncontrolled chambers are notably higher than in the controlled chamber confirming the effectiveness of the control strategy.

**Video S1. A) Movie of the *in silico* set-point control simulation**. (Left panel) Agent-based simulated set-point reference control experiment of bacterial cells in a microfluidic chamber; (top right panel) cell count over time; (bottom right panel) quantified GFP fluorescence (green) and input (IPTG in red) calculated by the relay controller. **B) Movie of the *in silico* time-varying control simulation.** (Left panel) Agent-based simulated time-varying reference control experiment of bacterial cells in a microfluidic chamber; (top right panel) cell count over time; (bottom right panel) quantified GFP fluorescence (green) and input (IPTG in red) calculated by the relay controller.

**Video S2 Movie of the experiment in Figure 3A.** (Top left panel) Segmentation mask calculated at every sampling time; (bottom left panel) bacteria cell fluorescence during the whole experiment; (top right panel) cell count over time; (bottom right panel) desired ($y_{ref}$ in blue) experimentally quantified GFP fluorescence ($y$ in green) and input (IPTG in red) calculated by the control algorithm are shown for the whole duration of the experiment.

**Video S3 Movie of the experiment in Figure 4A.** (Top left panel) Segmentation mask calculated at every sampling time; (bottom left panel) bacteria cell fluorescence during the whole experiment; (top right panel) cell count over time; (bottom right panel) desired ($y_{ref}$ in blue) experimentally quantified GFP fluorescence ($y$ in green) and input (IPTG in red) calculated by the control algorithm are shown for the whole duration of the experiment.
